# Landscape heterogeneity of peasant-managed agricultural matrices

**DOI:** 10.1101/668103

**Authors:** Ana L. Urrutia, Cecilia González-González, Emilio Mora Van Cauwelaert, Julieta A. Rosell, Luis García Barrios, Mariana Benítez

## Abstract

In agricultural landscapes, management practices and other environmental and social factors shape complex agroecological matrices. In turn, the structure of such matrices impacts both agricultural activities and biodiversity conservation, for instance, by mediating wildlife migration between agricultural and habitat patches. One way to characterize a matrix, its potential role in biodiversity conservation, and how its descriptors change across different spatial scales, is characterizing heterogeneity metrics and systematically examining how such metrics change with grain size and landscape extent. However, these methods have rarely been applied to tropical, peasant-managed landscapes, even though this type of landscape occupies most of the agricultural surface in or near biodiversity hotspots. We focus on a peasant-managed agricultural landscape in Oaxaca, Mexico, for which we mapped and quantified the land-use classes and evaluated heterogeneity metrics. We also examined the response of heterogeneity metrics to changes in grain and extent scales. This allowed us to further understand the structure and conservation potential of the agroecological matrix in this type of landscape, to broadly compare this landscape with other agricultural landscapes in North America, and to recommend specific landscape metrics for different types of studies involving agricultural matrices. We conclude that this type of agricultural matrix is ideal to pursue joint agricultural and conservation strategies in an integrated landscape.

## 1. INTRODUCTION

Biodiversity conservation research at the landscape scale has mainly focused on the characteristics of habitat patches (primary vegetation). Nevertheless, attention to the matrix that surrounds the habitat has increased in recent years (Franklin and Lindenmayer, 2009; Perfecto and Vandermeer, 2009; Fahrig et al., 2011). Several studies have shown that this matrix has a great impact on species persistence, preventing or allowing migration and re-colonization amongst habitat patches in the landscape and promoting or preventing regional extinctions (Levins, 1969; Dunning et al. 1992; Perfecto et al. 2009; Franklin and Lindenmayer, 2009; Tscharntke et al. 2012). It is also known that the matrix is not homogeneous nor is it equally beneficial for the persistence of all species. The quality of the matrix, understood as its permeability for the transit of local biodiversity, thus varies largely depending on human management (Vandermeer and Perfecto, 2007; Perfecto et al. 2009; Fahrig et al. 2011; Tscharntke et al. 2012).

Currently, most of the world’s habitat matrix composition is agricultural (Foley et al., 2005; Fahrig et al., 2011; Kremen, 2015). This highlights the role of agricultural practices and their distribution in a landscape on biodiversity conservation. The do-called agroecological matrix can act as a refuge for biodiversity, as a facilitator of movement of organisms among habitat patches, and may help maintain metapopulation dynamics and long-term survival of forest species (Vandermeer, 2007; Vandermeer & Carvajal, 2001; Philpott et al., 2008). However, the realization of these potential roles on biodiversity conservation largely depends on the type of agricultural practices and the spatial organization of agriculture and other land-use classes in the matrix (González-González et al., 2016; Ramos et al., 2018). Indeed, it has been suggested that highly heterogeneous landscapes dominated by small-scale or peasant agriculture integrate agricultural and conservation activities in a better way than landscapes dominated by large-scale monocultures (Perfectto et al., 2009; Fahrig et al., 2011; Kremen, 2015).

Landscape heterogeneity is associated to spatial patterns that drive ecological processes at several scales (Levin, 1992; Wu et al., 2002). Spatial heterogeneity is divided in to compositional heterogeneity, which refers to the number and types of patches that constitute a landscape, and configurational heterogeneity, which corresponds to the arrangement of these patches in the landscape, how fragmented the landscape is, the density of its borders and the connectivity among classes of patches, among other features (Wu et al., 2002; Wu, 2004; Fahrig, 2011).

Ecological patterns and processes that matter at a certain scale may not be relevant at another one. For this reason, a great deal of information can be potentially lost when delimiting a particular spatial area and a particular grain size to study a landscape. These scale-related issues have been studied for decades as part of the modifiable areal unit problem, and they have driven the sought for methods capable of preserving information across different scales or at least able to quantify the loss of information when scales change (Turner 1990; Wu, 2004; Zhang and Li, 2013; Teng et al., 2016). Metrics that characterize the heterogeneity of a landscape are scale-dependent, that is, they change with grain size and the extent of the area under study (Wu et al., 2002). To address how landscape metrics change with grain size or landscape area, several authors have used scalograms (Wu et al., 2002; Wu, 2004; Zhang and Li, 2013; Teng et al., 2016). Based on these graphic representations, metrics can have diverse behaviors that can be classified based on how consistent these behaviors are across landscapes, how predictable responses are to changes of scale, and whether relationships between scale and metric can be represented with simple scaling equations or are staircase-like or erratic (Wu et al., 2002; Wu, 2004; Zhang and Li, 2013; Teng et al., 2016). Examining how metrics behave with scale changes is thus crucial because the interpretation and the comparison of landscape metrics across different conditions and with different objectives could differ markedly if a metric is sensitive to scale changes. Therefore, in order to fully understand the ecological processes occurring in a landscape, it is crucial to explore how the inference of landscape properties change along scales (Wu et al., 2002; Peters et al., 2007; Wu, 2004).

Landscape studies examining the effect of scale change have been carried out mainly in simulated landscapes or in temperate regions (McGarigal and Cushman, 2002; Fahrig, 2003; Shen, 2004), which tend to have fewer land uses and land-cover types (Gliessman, 1992; Brown, 2014) then the tropics. Given that geological, biological, and human processes that determine these landscape properties, and thus landscape heterogeneity, can be very different in the tropics, studies examining landscape heterogeneity metrics are urgently needed for these areas of the world. Agricultural landscapes in the tropics tend to be characterized by small plots or farms managed by peasants. Tropical agricultural landscapes, are highly fragmented, highly diverse in terms of land use, and highly connected among land-use classes (Fahrig et al., 2011). Although heterogeneity studies have been conducted in tropical areas, they have mostly focused on fragmented tropical forests (Concepcion et al. 2008; Arroyo-Rodríguez et al. 2017; Sánchez-de-Jesús; 2015), and have not usually considered agricultural landscapes. Considering the key role of the agricultural matrix in the conservation of tropical biodiversity (Perfecto et al., 2009), studies examining heterogeneity in agricultural landscapes could guide integrative productive and conservation policies. To the best of our knowledge no studies of landscape heterogeneity across scales have been conducted on rural or peasant-managed Latin American landscapes, which have had very different human influences than landscapes studied so far and are often located within or near biodiversity hotspots (McGarigal and Cushman, 2002; Sánchez-de-Jesús, 2015; Fahrig et al., 2011).

In this work, we focused on an agricultural landscape in a rural region in Mexico. This region is dominated by peasant agriculture, a kind of agriculture represented by smallholder farmers, with family management that produces, at least partially, for self-consumption. Peasant agriculture is highly representative of national agricultural practices as it contributes with 25.5 % of national maize production and has the potential half of the population of Mexico (CEMDA, 2017; Bellon et al., 2018). These practices are usually linked to a highly fragmented landscape as most of the farms belong to smallholders, and are very diverse in terms of land use, and management (Fahrig et al., 2011; Bellon et al., 2018). Based on a landscape of peasent-managed agroecosystems and forests, we examined the behavior of landscape heterogeneity metrics with changes in grain size (affecting level of resolution), and extension (total studied area), based on a land-use and vegetation map. Given the characteristics of rural agricultural landscapes largely managed by peasants, we would predict high number of patches, high patch richness and high values in connectivity and intercalation metrics, such as the intercalation and juxtaposition index. Our results show that there is high landscape fragmentation with potential for good connectivity, highlighting important differences between the heterogeneity of this peasant-managed landscape and previously studied agricultural landscapes with more intensive agricultural practices. Moreover, our results reveal that the performance of some of the widely used metrics could change markedly across scales in highly fragmented landscapes like the one we studied. We postulate specific metrics and scales that could work well in such landscapes and that could be used in future studies aiming to identify high quality matrices in landscapes with these characteristics, which happen to characterize tropical areas with very high biodiversity.

## 2. METHODS AND STUDY SITE

### 2.1 Study site

La Villa de Zaachila, henceforth Zaachila, is a municipality in the Central Valleys of the state of Oaxaca, Mexico. It is located 17 km south of the city of Oaxaca, between degrees 96 ° 40 ’and 96 ° 47’ longitude and 16 ° 54 ’and 17 ° 05’ latitude, and has an extension of 81 km^2^ and an altitude of 1500 m a.s.l. With two mountain ranges in the east and west edges and a valley in the center, Zaachila has a semi-dry-semi-warm climate.

The history of landscape management of Zaachila begins with the Zapotecs, about 3500 years ago. Throughout its history, the municipality has had different management and land holding systems (Ruiz Medrano, 2011). Today, agricultural plots are mostly managed by peasants for family or local consumption. In contrast with most agricultural sites in many other countries, a large percentage of these plots represent communal land, the so-called *ejidos* (INEGI 2010; Mora Van Cauwelaert, 2017). Historically, Zaachila has been an important point for regional commerce, as its traditional market has existed since the time of the Zapotecs, and even now it gathers farmers and peasants from all the surrounding villages. The village and the market are also reservoirs of great culinary diversity that is often associated with local landraces and management practices (Mora Van Cauwelaert, 2017).

### 2.2 Vegetation and land use map

To create the map, we used a remote sensing image (Copernicus Sentinel-2 2017) from the dry season (May of 2016) and included five land-use classes (Urban, Grassland, Rainfed agriculture, Irrigated agriculture, and Forest) based on the land use and vegetation use chart 1: 250 000 (Series V, INEGI 2005) and the land use map made by CIEC for North America (Colditz et al. 2012). The polygon delimitation corresponds to the political boundaries of the Zaachila municipality, which in turn correspond to the organization of public databases and field-work permits.

In order to map and verify specific land uses, we visited Zaachila and its surroundings in September and October 2017 and took 200 verification points. The status of the vegetation and land use classes was assessed by visual inspection with the support of local informers. We georeferenced points using the GPS map 64 of Garmin (error range ~ 3m), with a minimum distance of 5m between each reference point.

We performed a supervised classification with the maximum likelihood algorithm (Chuvieco, 1996; Schuster et al., 2012). We based the selection of the training areas on the Normalized Vegetation Index (NDVI) and the combination of natural and infrared RGB bands. These areas included 404 verification points and an interactive process of composition of bands linked to two-dimensional dispersion graphics, which made it possible to differentiate the assigned categories. We determined the accuracy of the classification with 78 validation points, using the confusion matrix and the Cohen’s Kappa statistic. We added the classes of water and greenhouses in QGIS 2.18.7 (QGIS Development Team 2017), which were plotted using the INEGI topographic chart for water bodies and corroboration field trips. However, in the final heterogeneity analyses, we did not consider the greenhouse class due to its negligible area (Series V, INEGI 2005). We reduced the salt and pepper effect with a manual reclassification.

### 2.3 Spatial heterogeneity characterization and scalograms

We characterized the site’s heterogeneity by estimating several landscape metrics. We calculated eight metrics of spatial heterogeneity for the landscape level and eight metrics for the class level. Most of the metrics taken for landscape level are the same for class level. Only four metrics are only for landscape level (patch richness and Shannon’s diversity index) or for class level (total (area) class and Percentage of landscape), therefore there are ten different metrics (Table 1). We selected these metrics to include both compositional and configurational heterogeneity, and for their suitability to explore the agricultural matrix. We included metrics for area, density, edge, shape, isolation, proximity, interspersion and diversity (Table 1). We analyzed spatial heterogeneity using FRAGSTATS 4.2.1 (McGarigal, et al. 2012).

**Table 1.**
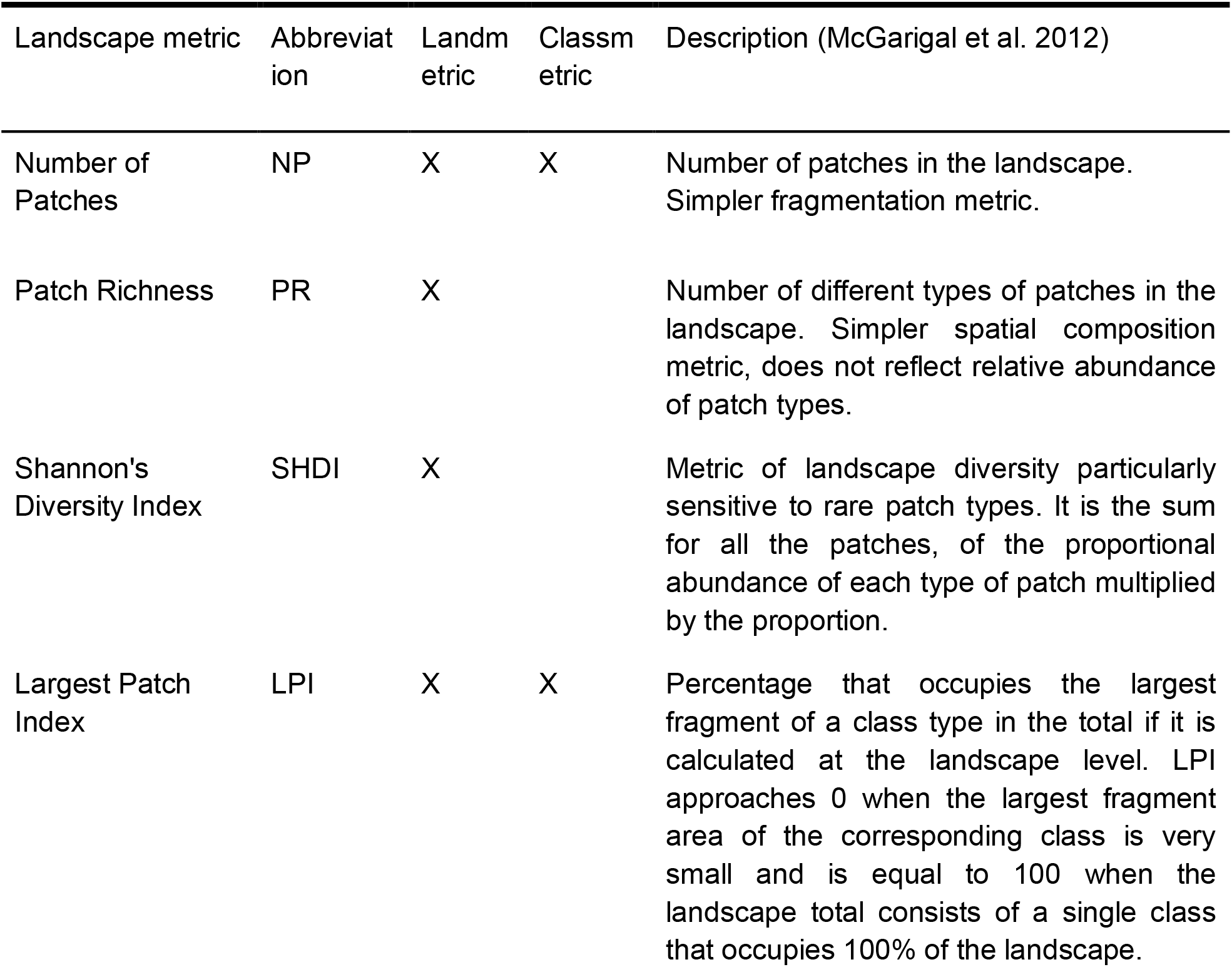
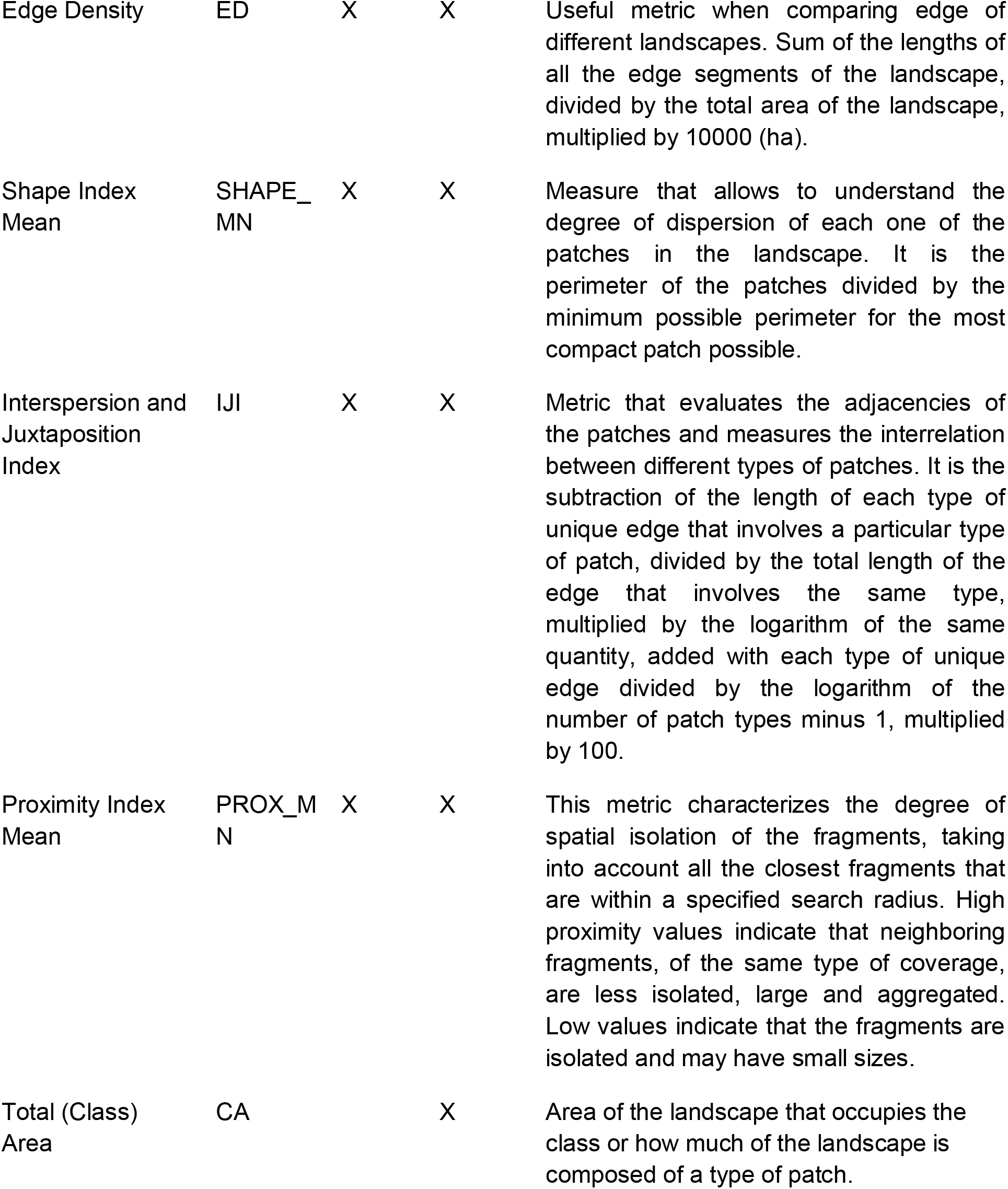
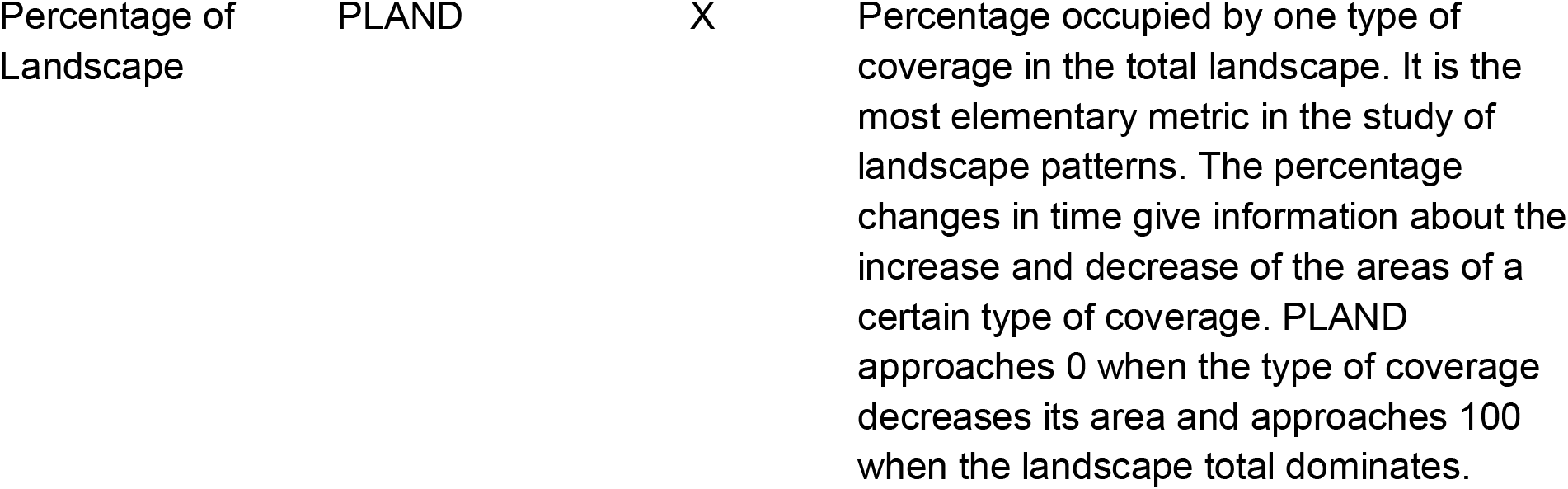
Landscape metrics used in this study. Most of the metrics calculated for landscape-level are the same for class level. Only four metrics are only for landscape level (PR and SHDI) or for class level (CA and PLAND).

In order to study the effect of the spatial scale on the landscape metrics, we built scalograms that described the response of each metric to changes in extension and grain size. Scalograms show how a metric responds to scale changes, and have been widely used to study sudden shifts in curves that could suggest hierarchical occurrences or critical scales (Zhang and Li, 2013). To test the effect of change in grain size we took the initial spatial resolution of the raster, which was 10 × 10 m in a total extension of 81 km^2^. The spatial resolution of the entire map was increased by 1 until it reached 100 × 100 pixels, keeping the extension constant. We assigned the grain identity resampling by majority. As for the effect of the extension, we took the complete extension of the quadrant surrounding the municipality (441 km2) and reduced it by 10 km2 until it reached 10 km2 (a similar approach to generate grain and extension scalograms was followed by Wu et al., 2002; Wu, 2004; Zhang and Li, 2013; Teng et al., 2016). Thus, we generated a total of 2000 images to represent the vegetation and land use of Zaachila.

We built scalograms plotting metric values against grain size or extension in R 3.4.3 (R Core Team, 2017). We used scalograms to: i) describe the behavior of the class level metrics, ii) classify the landscape level metric behavior in the face of scale changes, iii) test if changes in the metric with scale variation could be fitted with simple functional models, and iv) explore the sensitivity of metrics to scale change (Supplementary material Table 1, Table 2 and Table 3). Besides describing the overall trends of metric behavior with grain or extent scale change, it is important to characterize their sensitivity to small changes in the scale. To that end, we employed the coefficient of variation (CV), which expresses the standard deviation as a proportion of the average metric value. We used the CV to compare variation levels across landscape metrics. The larger the CV, the higher the sensitivity of the metric to changes in scale was (Teng et al., 2016) (Supplementary Material Table 3).

To examine whether a simple model could be fit to scalograms we used linear, polynomial or exponential functions and calculated residuals. Model residuals were used to visually inspect whether there were trends in the lack of fit with the change in grain size or extension (predictor variable in models) (Supplementary Material Figures 1 and 2).

### 2.4 Comparisons with other agricultural landscapes

Once landscape metrics were calculated, we compared them with those of 12 other agricultural landscapes of the METALAND database (Cardille et al., 2005). METALAND includes landscape metrics previously calculated with FRAGSTATS in landscapes of 6.5 km × 6.5 km in 8 million km2 in the USA. We chose 12 landscapes that had the same percentage of agricultural land use as Zaachila, and had similar percentages of urban land use and primary vegetation. For it to be comparable with the Zaachila landscape, we took a fragment of the map of the municipality that had an extension (6.5 km × 6.5 km) and grain size (30 × 30 m) equal to the one obtained in METALAND. We compared most of the landscape level metrics from Zaachila with the METALAND database, with the exception of proximity index mean and patch richness, metrics that were not comparable because of the particular characteristics that have to be defined for each metric.

## 3. RESULTS

### 3.1 Land use and vegetation characterization

The general accuracy of the land use map was 88.85 %, with a Kappa statistical index of 0.85, which is considered to be well above the minimum Kappa values required for reliable land use and vegetation maps (Congalton, 1991). The landscape of Zaachila is mainly agricultural (48 %), being rainfed agriculture the most prevalent land use class in the municipality (39 %). The next most represented class is the secondary forest (23 %), followed by the urban zone (19 %). The classes with less coverage in the landscape are water bodies and greenhouses (Figure 1). This makes Zaachila a predominantly rainfed agricultural landscape and a valuable site to study an agricultural matrix connecting habitat patches (Figure 1).

**Figure 1.**
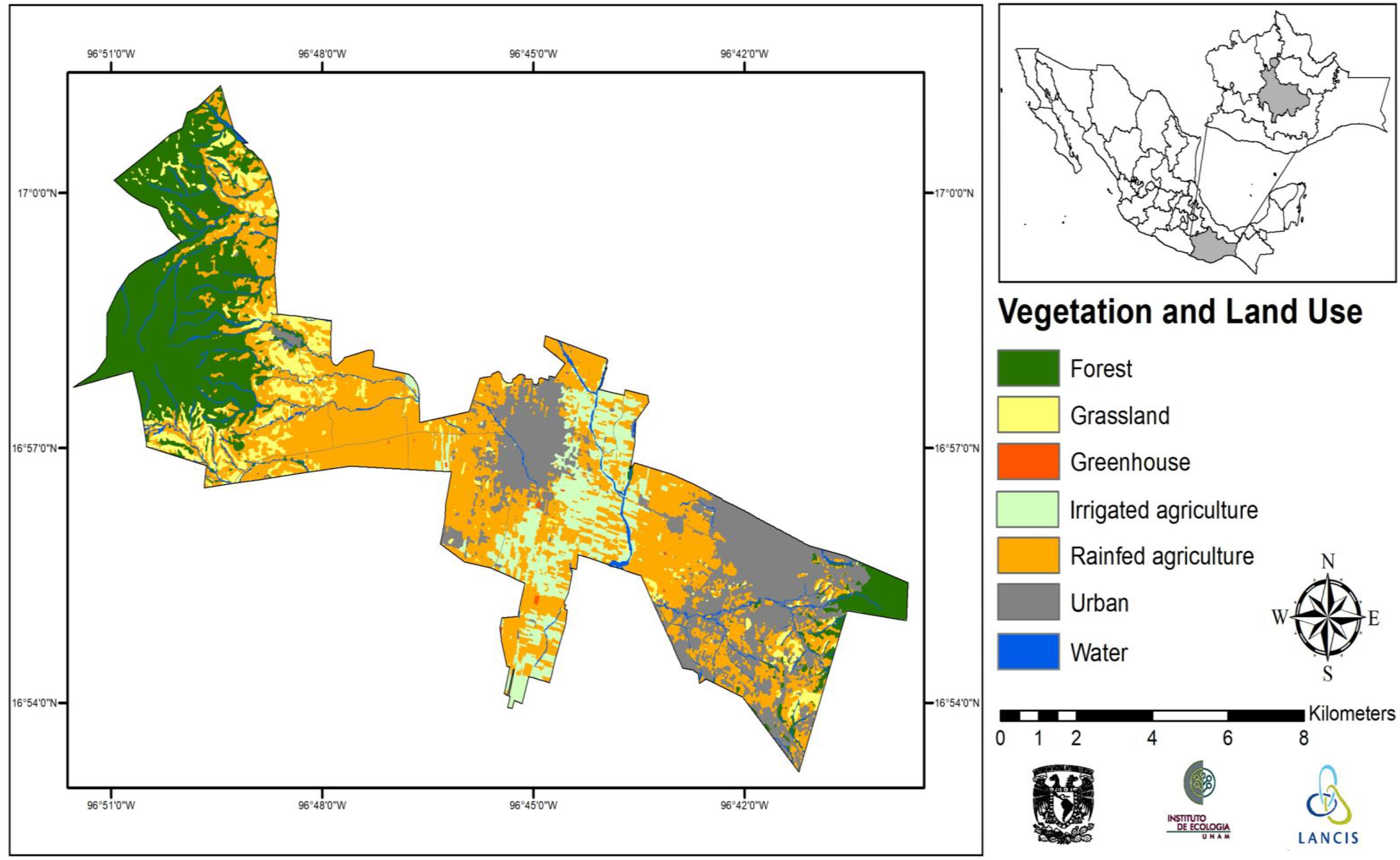
Zaachila’s vegetation and land use map with seven patch types, grain of 10 × 10 m and extent of 81 km2.

### 3.2 Landscape characterization

In terms of landscape level metrics, we found a very high number of patches (1867) and Shannon diversity index (1.54) (see also section below), with high dominance of forest patches in terms of area (the largest forest patch occupies 13.8 % of the landscape). The edge density was 125.13 m per ha and the average shape was 1.55, which means that the patches tend to be elongated. The interspersion and juxtaposition index was 74.97 % and the average proximity index mean within an area of 400 m was 1140, describing an intricate array of patches with a relatively high connectivity among them (Supplementary Material Table 4).

Regarding the class level metrics, the rainfed agriculture patches had the largest edge density (85.57 m/ha) and the highest interspersion and juxtaposition index (88.31 %), which means that this patch type is the most connected with other patch types, even if the largest patch in the landscape is a forest patch. However, the patch type with the highest proximity index mean (2961.07) was urban, which means that urban patches are the less isolated (Supplementary Material Table 5).

### 3.3 Response of metrics to changing scale at the landscape level

We plotted scalograms for the metrics in Table 1 and classified their behavior following Wu and collaborators (2002, 2004) and Teng and collaborators (2016) into four types: 1) consistent scaling relation (adjusted to a linear, polynomial or exponential function), 2) staircase-like response 3) invariant or non-answering to scale change and 4) erratic response (Figure 2, Table 2 and Supplementary Material Figure 1). In most cases, data violated homoscedasticity assumptions, so fits were only used for visual inspection of trends (Figure 2). Type 1 scalograms were the most common and pointed to metrics that are useful to describe matrix heterogeneity, as they appear to have a predictable behavior in response to scale changes. Metrics with type 1 scalograms were number of patches, largest patch index, shape index mean and interspersion and juxtaposition index (Figure 2 and Table 2).

**Figure 2.**
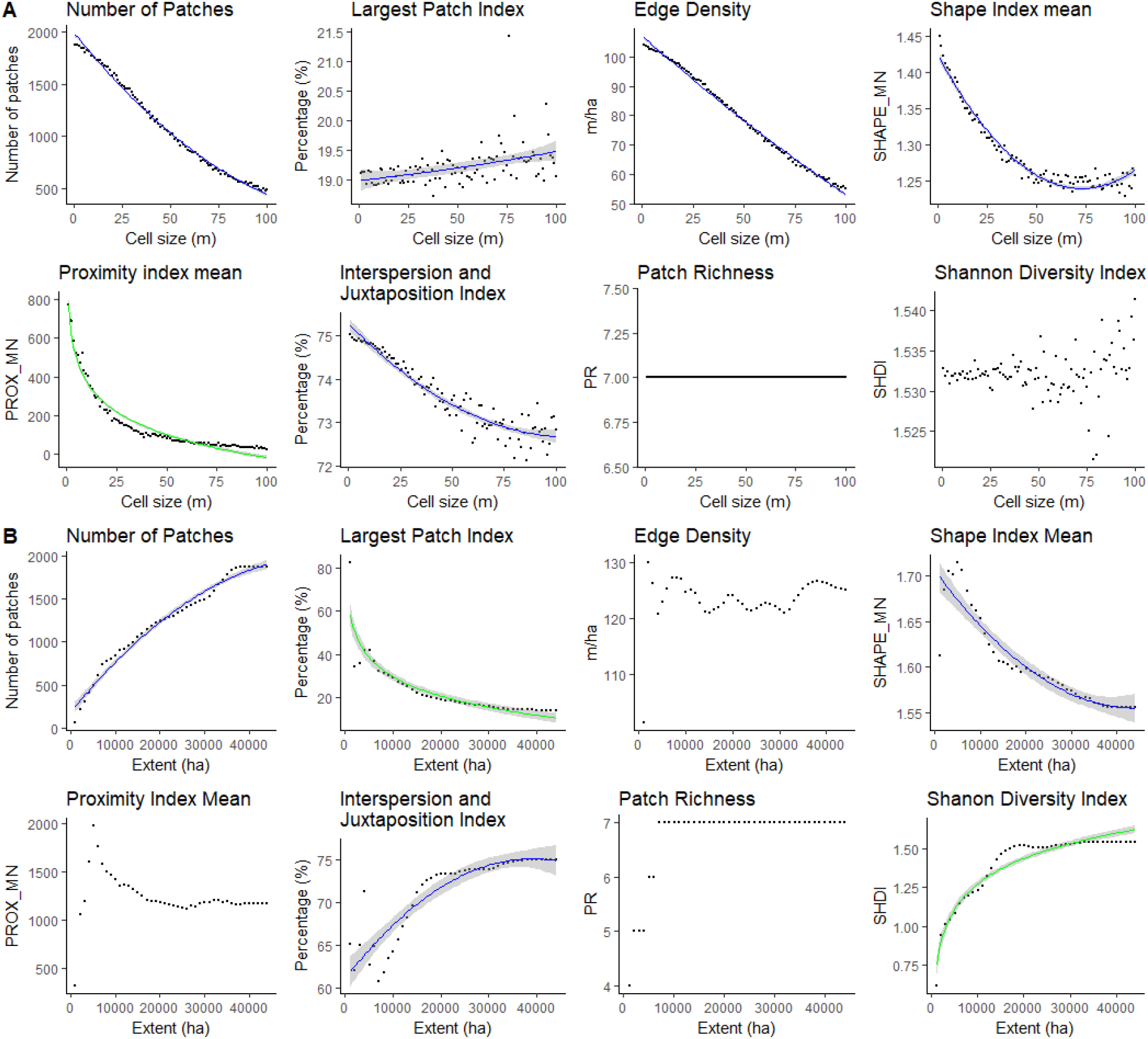
Scalograms showing the effects of changing grain size (a) and extent (b) on landscape metrics for Zaachila. Forscalograms with type 1 (consistent scaling relations) we present in green logarithmic fits and in blue power law fits.

**Table2.** Categories of behaviors observed in scalograms for landscape level

| Scale change | Metric | Response | Scaling relationship | Direction |
| --- | --- | --- | --- | --- |
| Grain size | NP | Type 1 | Power law | Decreasing |
| Grain size | LPI | Type 1 | Power law | Increasing |
| Grain size | ED | Type 1 | Power law | Decreasing |
| Grain size | SHAPE_MN | Type 1 | Power law | Decreasing |
| Grain size | PROX_MN | Type 1 | Logarithmic | Decreasing |
| Grain size | IJI | Type 1 | Power law | Decreasing |
| Grain size | PR | Type 3 | - | - |
| Grain size | SHDI | Type 4 | - | - |
| Extent | NP | Type 1 | Power law | Increasing |
| Extent | LPI | Type 1 | Logarithmic | Decreasing |
| Extent | ED | Type 4 | - | - |
| Extent | SHAPE_MN | Type 1 | Power law | Decreasing |
| Extent | PROX_MN | Type 4 | - | - |
| Extent | IJI | Type 1 | Power law | Increasing |
| Extent | PR | Type 2 | - | - |
| Extent | SHDI | Type 1 | Logarithmic | Increasing |

### 3.4 Responses of metrics to changing scale at the class level

We also plotted scalograms for class level metrics (Table 1). In general, the behavior of each class level metric was different, therefore, we could not classify them as we did for landscape level metrics. The scale change of the extension had a greater effect on these metrics than the change of grain size. For Zaachila, for an extension greater than 300 km^2^ we can observe a stabilization of the metrics; total (class) area, percentage of landscape, number of patches, largest patch index, edge density and interspersion and juxtaposition index (Figure 3B). For extension <300 km^2^, metrics had steps or non stabilized behavior. Moreover, for a couple of metrics (number of patches and proximity index mean), there seems to be a 60 m grain size threshold above which information for each class is lost (Figure 3A and Supplementary Material Figure 2).

**Figure 3.**
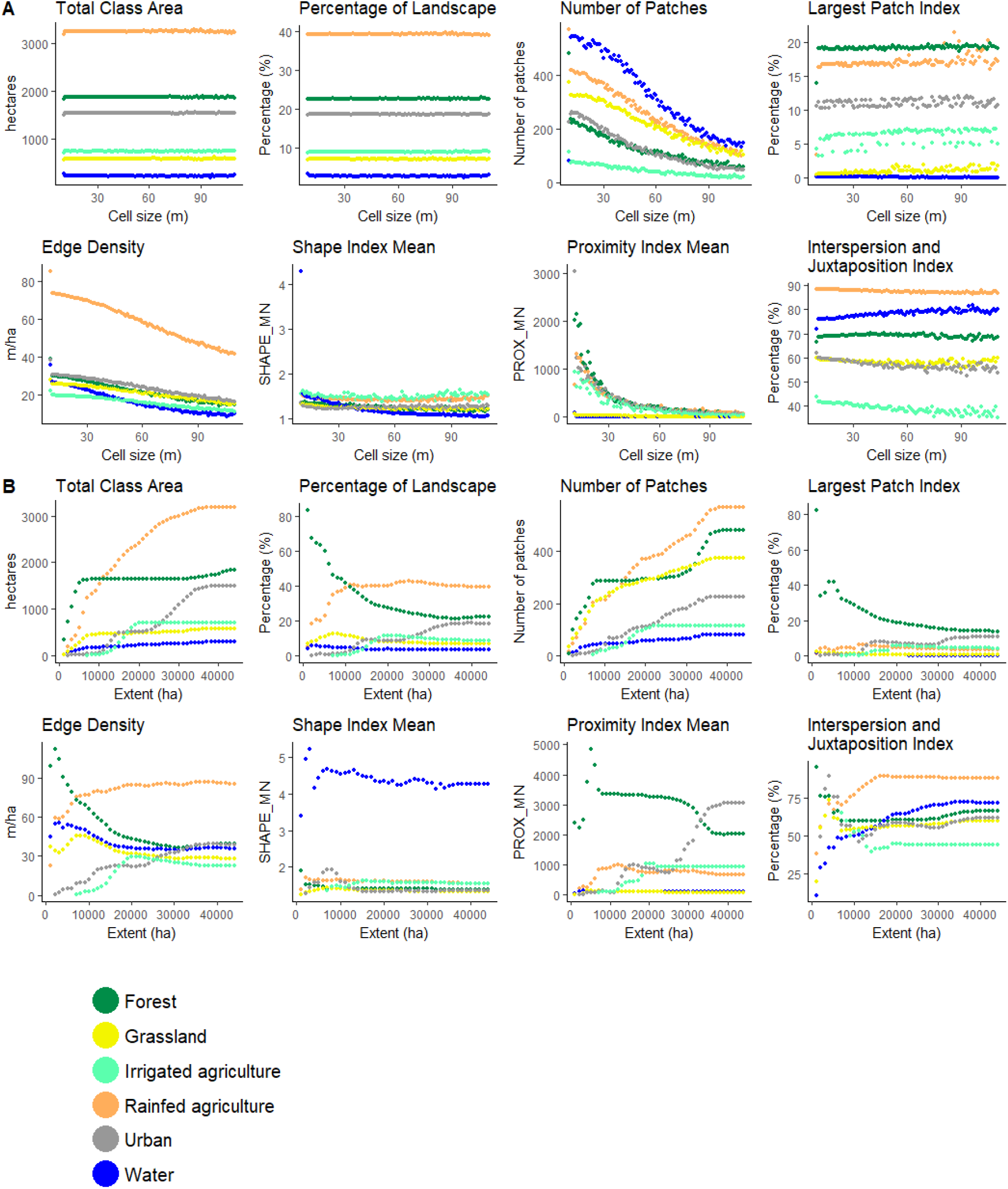
Scalograms showing the effects of changing grain size (a) and extent (b) on the class metrics taken for Zaachila’s vegetation and land use map. Since the behavior types for class level are highly different from class to class the classification was different and it is not possible to graphic the strongest scaling relations.

### 3.5 Comparison with metrics and scalograms in other agricultural systems

Compared with the values of the average metrics of twelve North American landscapes, the number of patches and the Shannon index in Zaachila were strikingly high. On the other hand, the shape index mean was visibly lower, perhaps due to the arrangement and distribution of small agricultural units into elongated plots, unlike the large land units of North American farmers (Figure 4). It is also worth noting that the Zaachila landscape exhibits a significantly higher patch diversity (Shannon index, Figure 4) and significantly less dominance of any class (Largest patch index, Figure 4) than agricultural landscapes in North America.

**Figure 4.**
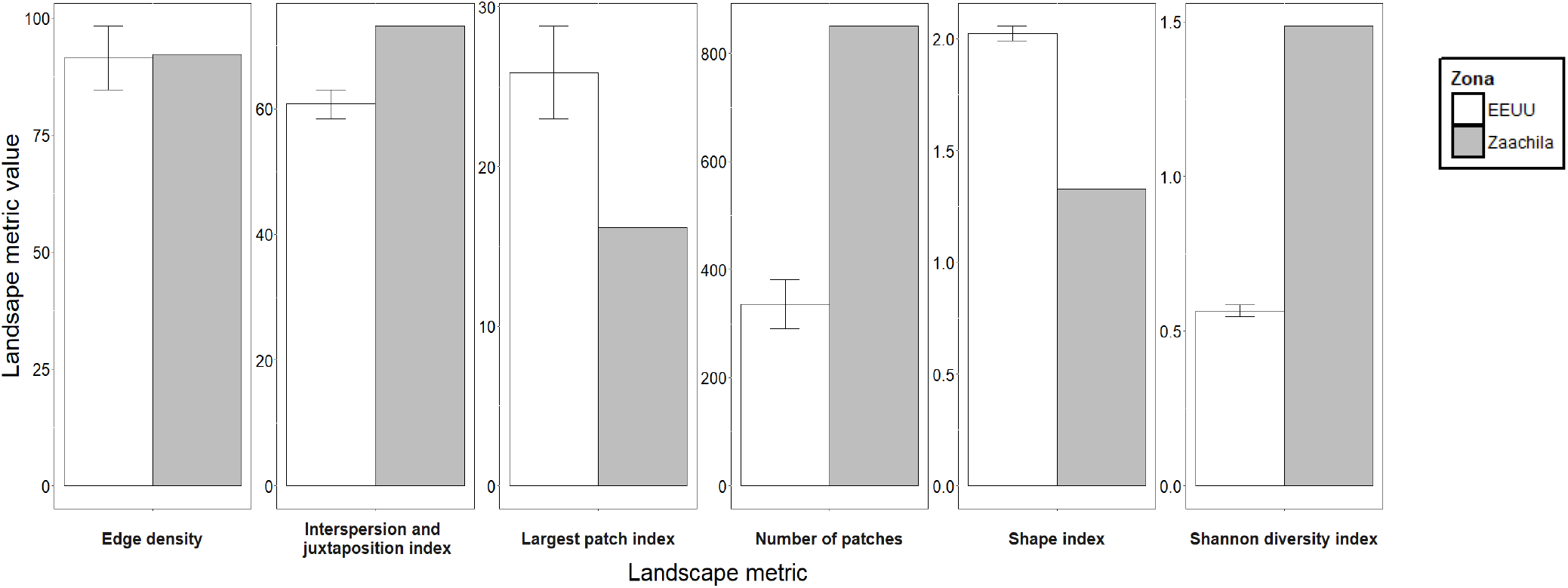
Bar graph comparing six landscape metrics from Zaachila versus thirteen different landscapes from the United States of America. The y axe is a compound of all the possible values of the different landscape metrics.

### 3.6 Sensitivity of the matrix descriptors response to scale change

Sensitivity can be understood as the size of variations or fluctuations around fitted models and can be examined through residuals (when it is possible to fit a functional model) and also using the coefficient of variation (see Methods). Most of the landscape level metrics show low sensitivity for small changes in scale. Class level metrics, on the contrary, show relatively high sensitivity for small changes in scale (Supplementary material Table 3).

## 5. DISCUSSION

In this work, we have characterized elements of the composition and configuration of a tropical, peasant-managed landscape in Oaxaca, Mexico, which is representative of largely understudied agricultural landscapes of the tropics, which are the ones with the largest potential to implement joint agricultural and conservation strategies.

Overall, Zaachila’s landscape is considerably fragmented with a high number of patches and a small largest patch index, especially when compared to North American landscapes of comparable size and agricultural use. Also, while the edge density between both types of landscapes was similar, the interspersion and juxtaposition index was higher in Zaachila (Figure 4), revealing an intricate arrangement of patches. Notably, the high patch interspersion and juxtaposition that characterizes Zaachila suggests that this landscape has large connectivity among patches. The Shannon index for patch diversity was also significantly higher in Zaachila in comparison with North American landscapes. Altogether, the diversity, complexity and potential connectivity of the Zaachila landscape reveals how different the tropical and peasant-managed landscapes can be in comparison with the agricultural landscapes that have been the focus of agricultural landscape heterogeneity studies so far. As hypothesized, Zaachila’s landscape is associated to an agricultural matrix with high number of patches, high patch richness and high values in connectivity and intercalation metrics that coincide with an ideal matrix heterogeneity aimed at land integration strategies for agricultural production and biodiversity conservation (Perfecto & Vandermeer, 2010; Fahrig, 2011).

In order to further discuss the quality and potential of Zaachila’s agricultural matrix in terms of biodiversity conservation, we will focus on the metrics describing the forest and rainfed agriculture classes, the two key classes in terms of strategies to integrate agricultural production and biodiversity conservation. Rainfed agriculture has a large interspersion and juxtaposition index, edge density and has little dominance (small largest patch index) (Supplementary Material Table 5). This is, rainfed agriculture is rather atomized and in contact with all types of patches. In contrast, the forest patch distribution is dominated by the largest patches, has little edge density and small interspersion and juxtaposition index (Supplementary Material Table 5). In other words, the forest class is concentrated in few, large patches with relatively little contact with the rest of the matrix. Considering that forest patches are in general apart from each other and that this may hinder metapopulation migration and recolonization dynamics, the matrix surrounding forest patches becomes central for the conservation of wild metapopulations. Since such matrix is dominated by rainfed agriculture, which in turn has a potentially high connectivity, it is crucial to maintain and foster agricultural practices that provide this class with a high permeability for local biodiversity (Vandermeer y Perfecto, 2007; Gonzalez-Gonzalez, 2019).

It is noteworthy that half of the scalograms exhibit similar behavior at the landscape level for Zaachila and for other agricultural landscapes previously studied (Wu et al., 2002; Wu, 2004; Teng, 2016; Zheng, 2014) (Supplementary Material Table 6). This is the case for number of patches, largest patch index and edge density for grain size change. Number of patches, edge density and patch richness also showed the same behaviors for the change in extent. These results point to a set of metrics that are likely to be robust and reliable for its use in a wide range of landscapes differing in geography, land use management and history of management, such as number of patches, largest patch index and edge density for grain size change and number of patches, edge density and patch richness for extent (Supplementary Material Table 6). In contrast, the other half of the landscape metrics exhibited qualitatively different behaviors between this study and others in response to scale change, such as shape index mean and Shannon’s diversity index. Most of the differences were between responses reported as erratic or staircase-like responses in other studies but that presented consistent scaling relations in Zaachila. The different behavior of this group of metrics highlight the importance of testing metric behavior in landscapes with diverse geophysical and social contexts that might lead to differences in the configuration of heterogeneity.

Taking into consideration the metrics and their behaviors in different scales we reckon that the most appropriate metrics for a study depend strongly on the goals of each study. For comparative research between landscapes with different geography and management history, in which metrics may not have been taken at the same scale, we propose to focus on metrics that are sensible enough to show differences across landscapes, but that are invariant to scale change (type 3). Metrics with consistent scaling relations (type 1) could be somewhat useful given that metric changes based on scale can be anticipated and that their sensitivity to scale change (CV and residuals) is small or moderate (Supplementary Material Table 3, Figure 1 and Figure 2). In order for these metrics to be useful, their behaviors must be robust, that is, must have shown similar behavior in landscapes with very different geography, geomorphology, land use management and history of management. If the metrics comply with these characteristics we can assume a simple relation between the metrics no matter the scale at which they were taken.

In this study, metrics that had these characteristics include the number of patches, interspersion and juxtaposition index and largest patch index. The last two have small CV values which also indicate that they are metrics with relatively low sensitivity to scale change (Supplementary Material Table 3). These three metrics are useful for describing landscape fragmentation and class dominance and differed between Zaachila and other landscapes (Figure 4). They had behaviors type 1 and 3 (Figures 2 and 3), i.e., behaviors that make these metrics robust across different agricultural landscapes (Wu et al., 2002; Wu, 2004; Teng et al., 2016) and had low sensitivity to scale change (Supplementary Material Table 3 and Table 6).

For studies aiming to describe the potential of a landscape for having a good quality matrix, we propose those metrics that reveal more information about connectivity, fragmentation per se, diversity and permeability of the classes (although information beyond the landscape metrics is necessary for the characterization of the matrix quality, especially for the permeability (Watling et al., 2011)). Metrics that best describe the matrix quality for Zaachila are percentage of landscape, edge density, and interspersion and juxtaposition index. Nevertheless, metrics that allow us to make replicable and more detailed studies in tropical landscapes are all the metrics typified as type 1 and type 3 with low or moderate sensitivity to scale change because they can be interpolated or extrapolated through the different landscapes and scales accurately using little information (Wu et al., 2002). Even within metrics with erratic behavior (type 4) there is a range of scales in which some metrics may be useful for describing different tropical, peasant-managed landscapes. Such is the case of Shannon’s diversity Index, that has a consistent scaling behavior between grain sizes of 10 × 10 m2 and 50 × 50 m2 and in any extension. Another type 4 metric is the proximity index mean, that in an extension larger than 300 km2 has consistent scaling behaviors (Figure 2).

In terms of which scale should be used in future works that address how landscape features affect agricultural plots (e.g. Connelly et al. 2015; Poveda et al. 2012; Avelino et al. 2012), we found no characteristic scale for the landscape level metrics. For the class level metrics a grain size below 60 m is recommended, while the change of extent shows a pattern in which the metrics stabilize for all classes from 300 km2 onwards. Stabilization may be very good for comparisons among different landscapes because it allows comparing them knowing that the metrics will be robust. However, for studies within a single landscape it is interesting to retain variation among classes, so the best scale for studies within Zaachila’s municipality and similar landscapes should be smaller than 300 km2. This also suggests that a change in hierarchical structure happens at 300 km2 in this landscape (Zhang and Li, 2013). However, the results obtained for changes in extent scale must be taken with care because they are highly dependent on the starting point of the smallest extent.

Finally, it is important to consider some of the limitations of this study. The shape of Zaachila’s polygon is one of them, given that it has an irregular shape. Some of the agricultural landscapes studied before are also defined by irregular polygons (Zhang and Li, 2013; Teng et al., 2016) but most are squared landscapes (Wu et al., 2002; Wu, 2004 and Cardille et al. 2017). This is particularly relevant for extension scalograms. It is also necessary to highlight that more than 150 other metrics have been developed to describe landscapes and that they would also need to be examined in terms of their stability with scale changes. Although it is true that many landscape metrics are highly correlated (Wu et al., 2002), it is still necessary to characterize the correlation among this large set of potential metrics in the type of landscape that we study here. Moreover, we think this kind of studies should also take into account the effects of different time scales on the metric behavior. In Zaachila the main land use is rainfed agriculture, which is modified by peasants and other actors along seasons, years and decades, so that the values of the landscape metrics could exhibit non-trivial dynamics along time points. Even in the absence of these more detailed temporal studies, here we show that Zaachila’s agricultural, fragmented, and tropical landscape with a matrix conformed mainly by rainfed agriculture holds a great potential for conservation if agricultural practices that increase the permeability of the matrix are maintained or propitiated.

## ACKNOWLEDGEMENTS

The authors thank Irene Ramos and members of *La Parcela* Laboratory for their valuable comments and suggestions. Juan Arias del Angel, Alexandre Beaupré, Maria de Guadalupe León, Diego Contreras, Tania Lara, Justino López Angel, Raymundo Aguilar and his family, members of *El Molote* collective and the *Regiduría de agricultura del Municipio de Villa de Zaachila* helped in the field trips and recognition of the state of vegetation and land use of the municipality. We also thank M. en C. Coral Eloisa Rangel for her help in the elaboration of the vegetation and land use map. Cecilia González González and Ana L. Urrutia acknowledge the graduate program “Posgrado en Ciencias Biológicas, Universidad Nacional Autónoma de México” and CONACyT scholarship (817256). This article covers part of the requirements to obtain the M.Sc. Degree in Biological Sciences (Ecology). M.Benítez acknowledges financial support from UNAM-DGAPA-PAPIIT (IN207819).

