## Supplementary material for "Landscape heterogeneity of peasant-managed agricultural matrices"

### Suppementary material

Table 1. Regressions to test consistent scaling relations in the full landscape of Zaachila with respect to grain size.

| Metric-level | Function | variable | sqr_err | slope | intercept | R2 | est_F | valrP |
|---|---|---|---|---|---|---|---|---|
| Landscape | Linear | NP | 4030.61057 | -15.377 | 1865.328 | 0.98 | 4936.36349 | 0 |
| Landscape | Linear | LPI | 0.33315586 | 0.008 | 18.772 | 0.136 | 15.5866235 | 0 |
| Landscape | Linear | ED | 4.68701541 | -0.552 | 106.766 | 0.982 | 5464.67404 | 0 |
| Landscape | Linear | SHAPE_MN | 0.00111813 | -0.002 | 1.372 | 0.68 | 210.345225 | 0 |
| Landscape | Linear | PROX_MN | 15338.3178 | -4.961 | 407.854 | 0.577 | 135.049104 | 0 |
| Landscape | Linear | IJI | 0.08735667 | -0.026 | 74.887 | 0.866 | 640.399213 | 0 |
| Landscape | Linear | PR | 1.49E-28 | 0 | 7 | 0.499 | 98.7318367 | 0.086 |
| Landscape | Linear | SHDI | 9.29E-06 | 0 | 1.532 | 0.005 | 0.45952707 | 0.499 |
| Landscape | Polynomial | NP | 1088.35216 | -4505.387 | 1096.495 | 0.995 | 9180.81739 | 0 |
| Landscape | Polynomial | LPI | 0.32313987 | 2.302 | 19.165 | 0.162 | 9.47250681 | 0 |

| | | | | | | | | |
|---|---|---|---|---|---|---|---|---|
| Landscape | Polynomial | ED | 3.87025049 | -161.649 | 79.181 | 0.985 | 3285.87739 | 0 |
| Landscape | Polynomial | SHAPE_MN | 0.00025891 | -0.49 | 1.289 | 0.926 | 612.231564 | 0 |
| Landscape | Polynomial | PROX_MN | 6111.36602 | -1453.71 | 159.782 | 0.831 | 241.741686 | 0 |
| Landscape | Polynomial | IJI | 0.05935008 | -7.555 | 73.598 | 0.909 | 489.659894 | 0 |
| Landscape | Polynomial | PR | 1.41E-28 | 0 | 7 | 0.501 | 49.2055906 | 0.08 |
| Landscape | Polynomial | SHDI | 8.22E-06 | 0.002 | 1.532 | 0.119 | 6.635649 | 0.475 |
| Class | Linear | CA | 1098362.02 | 0.011 | 1179.109 | 0 | 6.51E-05 | 0.994 |
| Class | Linear | PLAND | 161.04702 | 0 | 14.286 | 0 | 6.93E-13 | 1 |
| Class | Linear | NP | 13219.9963 | -2.197 | 288.442 | 0.237 | 218.72805 | 0 |
| Class | Linear | LPI | 56.1854772 | 0.007 | 7.357 | 0.001 | 0.55897171 | 0.455 |
| Class | Linear | ED | 278.582376 | -0.158 | 32.081 | 0.07 | 53.4472247 | 0 |
| Class | Linear | SHAPE_MN | 0.0356948 | -0.001 | 1.367 | 0.045 | 33.1480987 | 0 |

| | | | | | | | | |
|---|---|---|---|---|---|---|---|---|
| Class | Linear | PROX_MN | 68244.0155 | -5.119 | 475.733 | 0.246 | 230.081572 | 0 |
| Class | Linear | IJI | 235.731889 | -0.031 | 65.04 | 0.004 | 2.47997145 | 0.116 |
| Class | Polynomial | CA | 1098361.77 | 8.469 | 1179.764 | 0 | 0.00011216 | 0.994 |
| Class | Polynomial | PLAND | 161.04702 | 0 | 14.286 | 0 | 2.80E-12 | 1 |
| Class | Polynomial | NP | 13159.9502 | -1702.876 | 156.642 | 0.24 | 111.313298 | 0 |
| Class | Polynomial | LPI | 56.1763146 | 5.612 | 7.791 | 0.001 | 0.33654789 | 0.455 |
| Class | Polynomial | ED | 278.515701 | -122.195 | 22.623 | 0.071 | 26.7763622 | 0 |
| Class | Polynomial | SHAPE_MN | 0.03473353 | -1.089 | 1.283 | 0.071 | 26.7504397 | 0 |
| Class | Polynomial | PROX_MN | 59862.9441 | -3968.152 | 168.605 | 0.339 | 180.242494 | 0 |
| Class | Polynomial | IJI | 235.727472 | -24.213 | 63.166 | 0.004 | 1.24484644 | 0.116 |

Table 2. Regressions to test consistent scaling relations in the full landscape of Zaachila with respect to extent.

| Metric-level | Function | variable | sqr_err | slope | intercept | R2 | est_F | valrP |
|---|---|---|---|---|---|---|---|---|
| Landscape | Linear | NP | 10839.9816 | 0.038 | 385.87 | 0.956 | 904.391746 | 0 |
| Landscape | Linear | LPI | 61.4125117 | -0.001 | 38.857 | 0.581 | 58.3425279 | 0 |
| Landscape | Linear | ED | 15.0570482 | 0 | 121.854 | 0.051 | 2.24302846 | 0.142 |
| Landscape | Linear | SHAPE_MN | 0.00043293 | 0 | 1.679 | 0.805 | 173.108686 | 0 |
| Landscape | Linear | PROX_MN | 47354.644 | -0.005 | 1360.553 | 0.089 | 4.09849898 | 0.049 |
| Landscape | Linear | IJI | 5.50221805 | 0 | 64.277 | 0.727 | 112.072756 | 0 |
| Landscape | Linear | PR | 0.31146614 | 0 | 6.064 | 0.323 | 20.0598611 | 0 |
| Landscape | Linear | SHDI | 0.01505598 | 0 | 1.104 | 0.652 | 78.8128688 | 0 |
| Landscape | Polynomial | NP | 4462.59949 | 3204.751 | 1245.727 | 0.982 | 1101.56018 | 0 |
| Landscape | Polynomial | LPI | 35.1435429 | -61.266 | 22.418 | 0.76 | 65.0856447 | 0 |
| Landscape | Polynomi | ED | 14.973690 | 5.948 | 123.45 | 0.0 | 1.2150283 | 0.1 |

| | | | | | | | | |
|---|---|---|---|---|---|---|---|---|
| | al | | 6 | | | 56 | 7 | 45 |
| Landscape | Polynomial | SHAPE_MN | 0.00031017 | -0.28 | 1.604 | 0.86 | 126.046243 | 0 |
| Landscape | Polynomial | PROX_MN | 47350.3009 | -450.916 | 1239.569 | 0.089 | 2.00252163 | 0.052 |
| Landscape | Polynomial | IJI | 3.77762978 | 25.417 | 71.096 | 0.813 | 89.0339696 | 0 |
| Landscape | Polynomial | PR | 0.15887438 | 2.558 | 6.75 | 0.655 | 38.8843944 | 0 |
| Landscape | Polynomial | SHDI | 0.00351584 | 1.115 | 1.403 | 0.919 | 232.021475 | 0 |
| Class | Linear | CA | 714420.419 | 0.021 | 367.874 | 0.085 | 27.5018214 | 0 |
| Class | Linear | PLAND | 236.20641 | 0 | 16.46 | 0.003 | 0.94703638 | 0.331 |
| Class | Linear | NP | 21018.3894 | 0.005 | 65.924 | 0.158 | 55.318844 | 0 |
| Class | Linear | LPI | 77.0364256 | 0 | 7.684 | 0.016 | 4.79475775 | 0.029 |
| Class | Linear | ED | 662.533324 | 0 | 40.124 | 0.005 | 1.56817779 | 0.211 |
| Class | Linear | SHAPE_MN | 1.13822735 | 0 | 1.976 | 0.003 | 0.99332462 | 0.32 |
| Class | Linear | PROX_MN | 1206330.8 | 0.008 | 663.85 | 0.0 | 2.4612488 | 0.1 |

| | | | | | | | | |
|---|---|---|---|---|---|---|---|---|
| | | | 4 | | 8 | 08 | 2 | 18 |
| Class | Linear | IJI | 196.628093 | 0 | 54.808 | 0.051 | 15.7267937 | 0 |
| Class | Polynomial | CA | 710631.721 | 4447.591 | 855.526 | 0.09 | 14.5610844 | 0 |
| Class | Polynomial | PLAND | 235.248723 | -15.007 | 14.815 | 0.007 | 1.0722647 | 0.331 |
| Class | Polynomial | NP | 20961.4804 | 1081.941 | 184.552 | 0.16 | 28.0395952 | 0 |
| Class | Polynomial | LPI | 76.1106583 | -19.284 | 5.57 | 0.028 | 4.20633898 | 0.029 |
| Class | Polynomial | ED | 656.395758 | -32.342 | 36.578 | 0.015 | 2.16324717 | 0.21 |
| Class | Polynomial | SHAPE_MN | 1.13669969 | -1.067 | 1.859 | 0.005 | 0.6932025 | 0.32 |
| Class | Polynomial | PROX_MN | 1206168.33 | 1728.933 | 853.425 | 0.008 | 1.24642381 | 0.118 |
| Class | Polynomial | IJI | 196.314147 | 55.797 | 60.926 | 0.052 | 8.08435682 | 0 |

Table 3. Sensitivity to scale change for the selected metrics

| **Metric level** | **Scale change** | **Metric** | **mean** | **sd** | **cv** | **Sensitivity** |
|---|---|---|---|---|---|---|

| | | | | | | |
|---|---|---|---|---|---|---|
| Landscape | Grain size | NP | 1096.495 | 455.034 | 41.499 | Moderate sensitivity |
| Landscape | Grain size | LPI | 19.165 | 0.624 | 3.256 | Low sensitivity |
| Landscape | Grain size | ED | 79.181 | 16.311 | 20.600 | Low sensitivity |
| Landscape | Grain size | SHAPE_MN | 1.289 | 0.059 | 4.577 | Low sensitivity |
| Landscape | Grain size | PROX_MN | 159.782 | 191.375 | 119.773 | High sensitivity |
| Landscape | Grain size | IJI | 73.598 | 0.812 | 1.103 | Low sensitivity |
| Landscape | Grain size | PR | 7.000 | 0.000 | 0.000 | Insensitive |
| Landscape | Grain size | SHDI | 1.532 | 0.003 | 0.196 | Insensitive |
| Landscape | Extent | NP | 1245.727 | 499.939 | 40.132 | Moderate sensitivity |
| Landscape | Extent | LPI | 22.418 | 12.253 | 54.657 | High sensitivity |
| Landscape | Extent | ED | 123.450 | 4.029 | 3.264 | Low sensitivity |
| Landscape | Extent | SHAPE_MN | 1.604 | 0.048 | 2.993 | Low sensitivity |
| Landscape | Extent | PROX_MN | 1239.569 | 230.617 | 18.605 | Moderate sensitivity |

| | | | | | | |
|---|---|---|---|---|---|---|
| Landscape | Extent | IJI | 71.096 | 4.545 | 6.393 | Low sensitivity |
| Landscape | Extent | PR | 6.750 | 0.686 | 10.163 | Moderate sensitivity |
| Landscape | Extent | SHDI | 1.403 | 0.211 | 15.039 | Moderate sensitivity |
| Class | Grain size | NP | 156.642 | 131.704 | 84.080 | High sensitivity |
| Class | Grain size | LPI | 7.791 | 7.504 | 96.316 | High sensitivity |
| Class | Grain size | ED | 22.623 | 17.324 | 76.577 | High sensitivity |
| Class | Grain size | SHAPE_MN | 1.283 | 0.193 | 15.043 | Low sensitivity |
| Class | Grain size | PROX_MN | 168.605 | 301.072 | 178.566 | High sensitivity |
| Class | Grain size | IJI | 63.166 | 15.391 | 24.366 | Low sensitivity |
| Class | Grain size | CA | 1179.764 | 1048.770 | 88.897 | High sensitivity |
| Class | Grain size | PLAND | 14.286 | 12.699 | 88.891 | High sensitivity |
| Class | Extent | NP | 184.552 | 158.253 | 85.750 | High sensitivity |
| Class | Extent | LPI | 5.570 | 8.863 | 159.120 | High sensitivity |
| Class | Extent | ED | 36.578 | 25.852 | 70.676 | High sensitivity |

| | | | | | | |
|---|---|---|---|---|---|---|
| Class | Extent | SHAPE_MN | 1.859 | 1.070 | 57.558 | High sensitivity |
| Class | Extent | PROX_MN | 853.425 | 1104.765 | 129.451 | High sensitivity |
| Class | Extent | IJI | 60.926 | 14.416 | 23.661 | Low sensitivity |
| Class | Extent | CA | 855.526 | 885.247 | 103.474 | High sensitivity |
| Class | Extent | PLAND | 14.815 | 15.420 | 104.084 | High sensitivity |

Table 4. Values obtained for Zaachila's landscape-level metrics at grain size 10 and for the whole landscape

| **Landscape metric (Landscape-level)** | **Result** |
|---|---|
| Number of patches(NP) | 1867 |
| Patch richness (PR) | 7 |
| Shannon's Diversity Index (SHDI) | 1.542 |
| Largest Patch Index (LPI) | 13.88% |
| Edge Density (ED) | 125.13 m/ha |
| Shape Index Mean (SHAPE_MN) | 1.56 |
| Interspersion and Juxtaposition Index (IJI) | 74.98 % |
| Proximity Index Mean (PROX_MN) | 1140.07 |

Table 5. Values obtained for Zaachila's class-level metrics at grain size 10 and for the whole landscape

| **Landscape metric (Class-level)** | **Urban** | **Irrigated Agriculture** | **Grassland** | **Rainfed Agriculture** | **Forest** | **Water** |
|---|---|---|---|---|---|---|
| Number of patches (NP) | 227 | 116 | 376 | 570 | 482 | 81 |
| Total (Class) Area (CA) | 1505.37 ha | 713.58 ha | 571.03 ha | 3187.98 ha | 1834.42 ha | 291.89 ha |
| Percentage of Landscape (PLAND) | 18.5585 % | 8.7972 % | 7.0398 % | 39.302 % | 22.6151 % | 3.5985 % |
| Largest Patch Index (LPI) | 10.5752 % | 4.3081 % | 0.5031 % | 3.4891 % | 13.8825 % | 0.2749 % |
| Edge Density (ED) | 38.68 m/ha | 22.39 m/ha | 28.12 m/ha | 85.57 m/ha | 39.05 m/ha | 35.90 m/ha |
| Shape Index Mean (SHAPE_MN) | 1.3597 | 1.5567 | 1.3341 | 1.5578 | 1.3728 | 4.2802 |
| Interspersion and Juxtaposition Index (IJI) | 61.89 % | 44.04 % | 59.85 % | 88.31 % | 66.52 % | 71.98 % |
| Proximity Index Mean (PROX_MN) | 2961.07 | 914.60 | 67.31 | 650.85 | 1965.03 | 84.18 |

Table 6. Comparison between the results obtained for Zaachila and landscapes studied by Wu et al.,(2002). (-) Is data not available.

| **Scale change** | **Metric** | **Response** | **Scaling relation** | **Direction** | **Wu** | **Scaling relation and direction** |
|---|---|---|---|---|---|---|
| Grain size | NP | Type 1 | Power law | Decreasing | Type 1 | Power function (D) |
| Grain size | LPI | Type 1 | Power law | Increasing | Type 1 | Power law or logatithmic (I) |

| | | | | | | |
|---|---|---|---|---|---|---|
| Grain size | ED | Type 1 | Power law | Decreasing | Type 1 | Power function (D) |
| Grain size | SHAPE_MN | Type 1 | Power law | Decreasing | Type 4 | |
| Grain size | PROX_MN | Type 1 | Logarithmic | Decreasing | Not available | |
| Grain size | IJI | Type 1 | Power law | Decreasing | Not available | |
| Grain size | PR | Type 3 | - | - | Type 2 | |
| Grain size | SHDI | Type 4 | - | - | Type 2 | |
| Extent | NP | Type 1 | Power law | Increasing | Type 1 | Power function (I) |
| Extent | LPI | Type 1 | Logarithmic | Decreasing | Type 4 | |
| Extent | ED | Type 4 | - | - | Type 4 | |
| Extent | SHAPE_MN | Type 1 | Power law | Decreasing | Type 4 | |
| Extent | PROX_MN | Type 4 | - | - | Not available | |
| Extent | IJI | Type 1 | Power law | Increasing | Not available | |
| Extent | PR | Type 2 | - | - | Type 2 | |
| Extent | SHDI | Type 1 | Logarithmic | Increasing | Type 2 | |

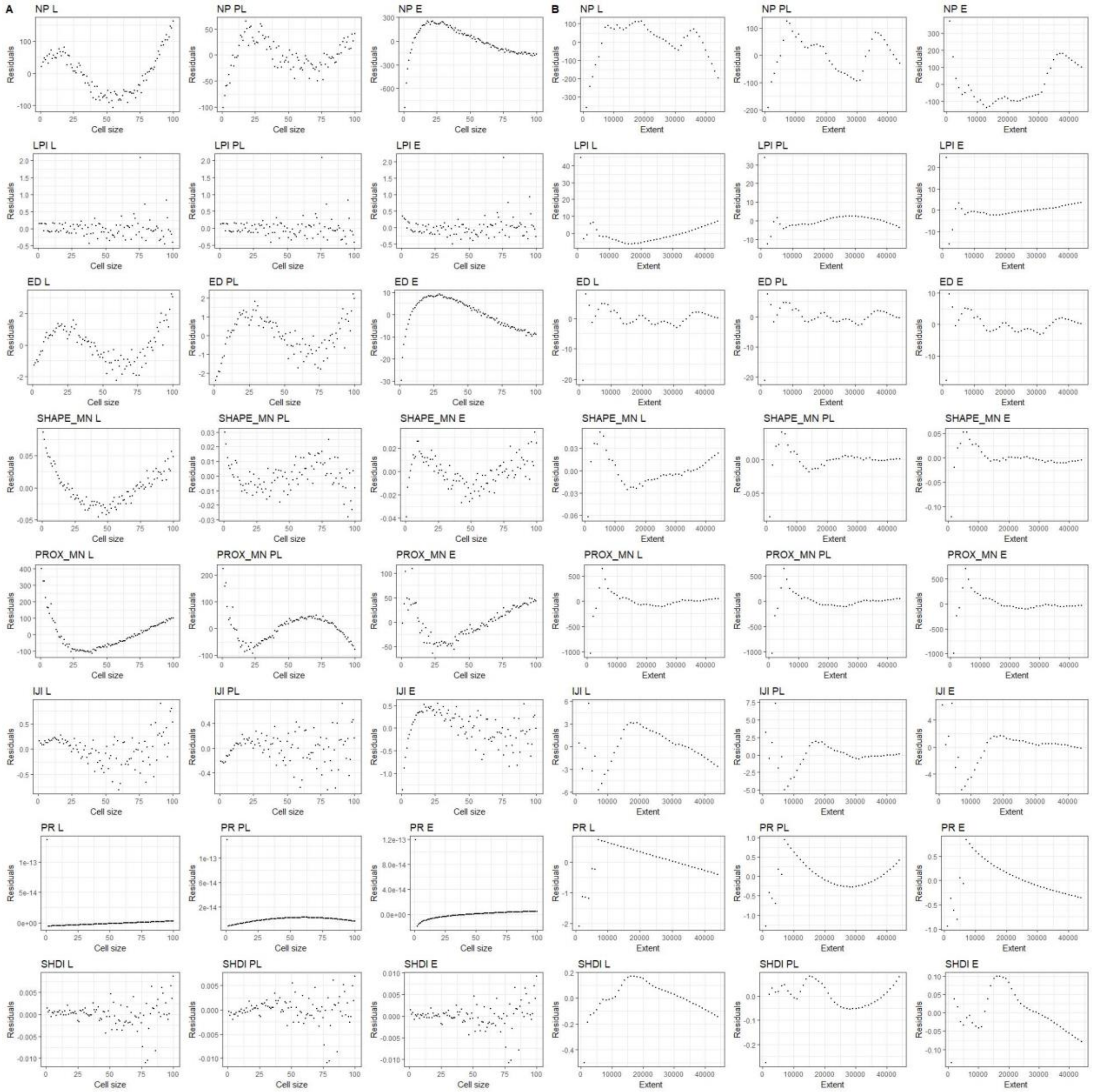

Figure 1. Landscape level metrics sensitivity tests. A) are the residuals values in different cell sizes and B) are the residuals in different extents. L stands for the sensitivity of the metrics to linear regressions, PL stands for the sensitivity of the metrics to polynomial regressions and E is for the sensitivity of the metrics to exponential regressions.

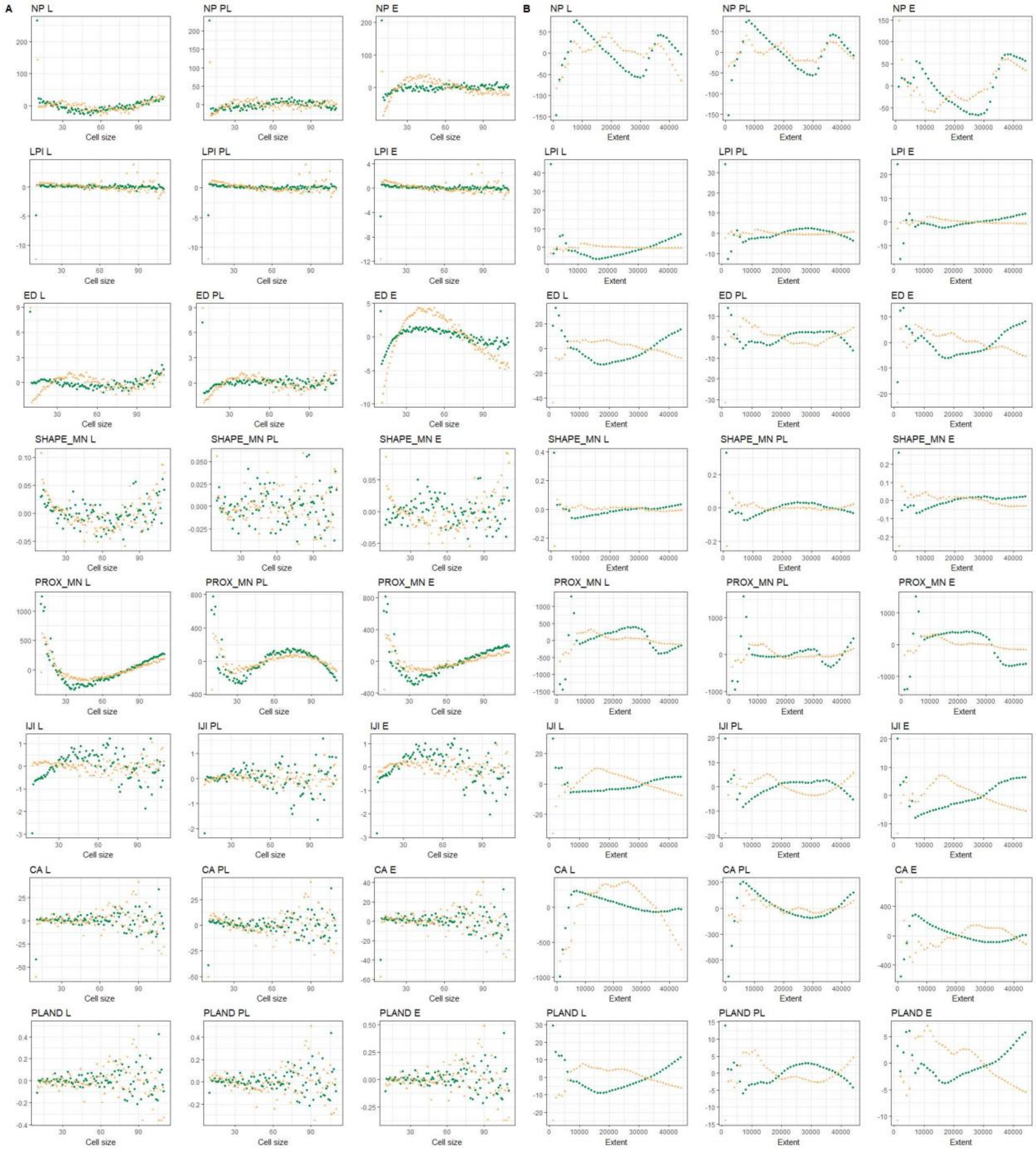

Figure 2. Class level metrics sensitivity tests for forest (green) and rainfed agriculture (yellow). A) are the residuals values in different cell sizes and B) are the residuals in different extents. L stands for the sensitivity of the metrics to linear regressions, PL stands for the sensitivity of the metrics to polynomial regressions and E is for the sensitivity of the metrics to exponential regressions.
